# Construction of a CMV promoter-driven Langat virus cDNA clone and reporter viruses

**DOI:** 10.1101/2025.11.13.688285

**Authors:** Anna I. Hermacinski, R. Blake Richardson, Eva Bednarski, Madihah Salim, Amit Garg, Jean K. Lim, Matthew J. Evans

**Affiliations:** Department of Microbiology, Icahn School of Medicine at Mount Sinai, New York, NY, 10029

**Keywords:** Langat virus, Flavivirus, Reverse genetics, Infectious clone, CMV promoter, Introns, Reporter virus

## Abstract

Langat virus (LGTV) is a tick-borne member of the *Flaviviridae* family and a biosafety level 2 surrogate for studying tick-borne encephalitis virus (TBEV) replication and pathogenesis. Here, we report the construction of a plasmid encoding a cytomegalovirus (CMV) promoter-driven LGTV cDNA that initiates infection following direct transfection of mammalian cells. Incorporation of three introns eliminated viral cDNA-associated toxicity in bacteria, enabling stable propagation of the full-length plasmid. Transfection of this construct resulted in high-level production of infectious LGTV, which exhibited robust replication kinetics, though slightly slower growth compared to a patient-derived isolate. We further engineered mCherry and Gaussia luciferase reporter versions of the clone, which yielded viruses expressing high levels of their respective reporters while retaining efficient replication. These LGTV infectious clones provide versatile tools for investigating viral replication, gene function, and pathogenesis, and may facilitate screening for antiviral inhibitors.

**Highlights:**

- Single-plasmid CMV-driven system launches infectious Langat virus (LGTV)
- Three introns stabilize full-length LGTV cDNA in E. coli
- DNA transfection yields high-titer infectious virus in mammalian cells
- Reporter LGTVs enable fluorescent and luminescent infection readouts
- Provides a versatile platform for LGTV genetics and pathogenesis studies

## Introduction

Langat virus (LGTV) is a tick-borne member of the family *Flaviviridae* in the genus *Orthoflavivirus* (Kwasnik et al., 2023). Like other flaviviruses, LGTV is a lipid-enveloped, positive-sense RNA arbovirus, but it is unique in that it is transmitted by ixodid ticks rather than mosquitoes. LGTV belongs to the tick-borne encephalitis virus (TBEV) serocomplex, which also includes Powassan and Louping-ill viruses. Although LGTV shares up to 84% amino-acid identity with TBEV, it is markedly less pathogenic (Pletnev and Men, 1998; Price et al., 1970). Whereas TBEV infection can cause severe neuroinvasive disease such as meningoencephalitis, LGTV infection is typically mild or subclinical and can be handled under biosafety level-2 (BSL-2) conditions. Owing to these characteristics, LGTV serves as a safe and tractable surrogate model for studying TBEV replication, immune evasion, and pathogenesis. The prototype LGTV TP21 strain was originally isolated from *Ixodes granulatus* ticks in Malaysia (Smith, 1956), whereas the E5 strain is a derivative obtained after approximately 42 serial passages of TP21 in embryonated chicken eggs (Thind and Price, 1966a, 1966b). Compared with TP21, the E5 strain is more attenuated in peripheral replication yet paradoxically exhibits enhanced neurovirulence and neuroinvasiveness in mice and nonhuman primates, likely reflecting adaptive changes acquired during egg passage (Price et al., 1970; Price and Thind, 1973).

Developing improved molecular tools for studying LGTV offers an opportunity to elucidate mechanisms that govern tick-borne flavivirus replication, immune evasion, and neurovirulence. Notably, LGTV served as the basis for an early live-attenuated TBEV vaccine developed in the 1970s but was later withdrawn from use (Price et al., 1970). Renewed study of LGTV may therefore inform the design of next-generation vaccines and antivirals as tick-borne encephalitis continues to rise in incidence and expand in geographic range across Europe and Asia (Kwasnik et al., 2023). Defining the viral determinants of infection, replication, and pathogenesis, and refining naturally attenuated strains for vaccine development, requires the ability to genetically manipulate viral genomes. Moreover, the incorporation of reporter genes to facilitate infection studies depends on robust reverse-genetics systems that permit precise control over viral sequence and phenotype.

Like other flaviviruses, LGTV encodes a single open reading frame that is translated into a polyprotein and co- and post-translationally cleaved into three structural proteins, capsid (C), precursor membrane (prM), and envelope (E), as well as seven nonstructural proteins, NS1, NS2A, NS2B, NS3, NS4A, NS4B, and NS5 (Fig. 1A) (Lindenbach et al., 2020). The structural proteins mediate viral entry and assembly, whereas the nonstructural proteins direct RNA replication and modulate host antiviral signaling pathways (Park et al., 2007). For most positive-strand RNA viruses whose genomes directly encode the proteins required for replication, transfection of a permissive cell with viral RNA is sufficient to initiate infection and produce progeny virions (reviewed in Aubry et al., 2015; Lindenbach, 2022). Such genomes can be cloned into plasmids containing promoter and terminator elements that enable *in vitro* transcription of infectious RNA for transfection. Because orthoflaviviruses possess capped RNA genomes, this transcription step can be circumvented by transfecting a plasmid that encodes the viral genome under the control of a polymerase II promoter, with the authentic 3’ terminus generated by a hepatitis D virus ribozyme (HDVr). In this system, the plasmid serves only to initiate the first round of RNA transcription, after which the viral RNA functions as both the translation and replication template to establish infection in transfected cells.

**Figure 1.**
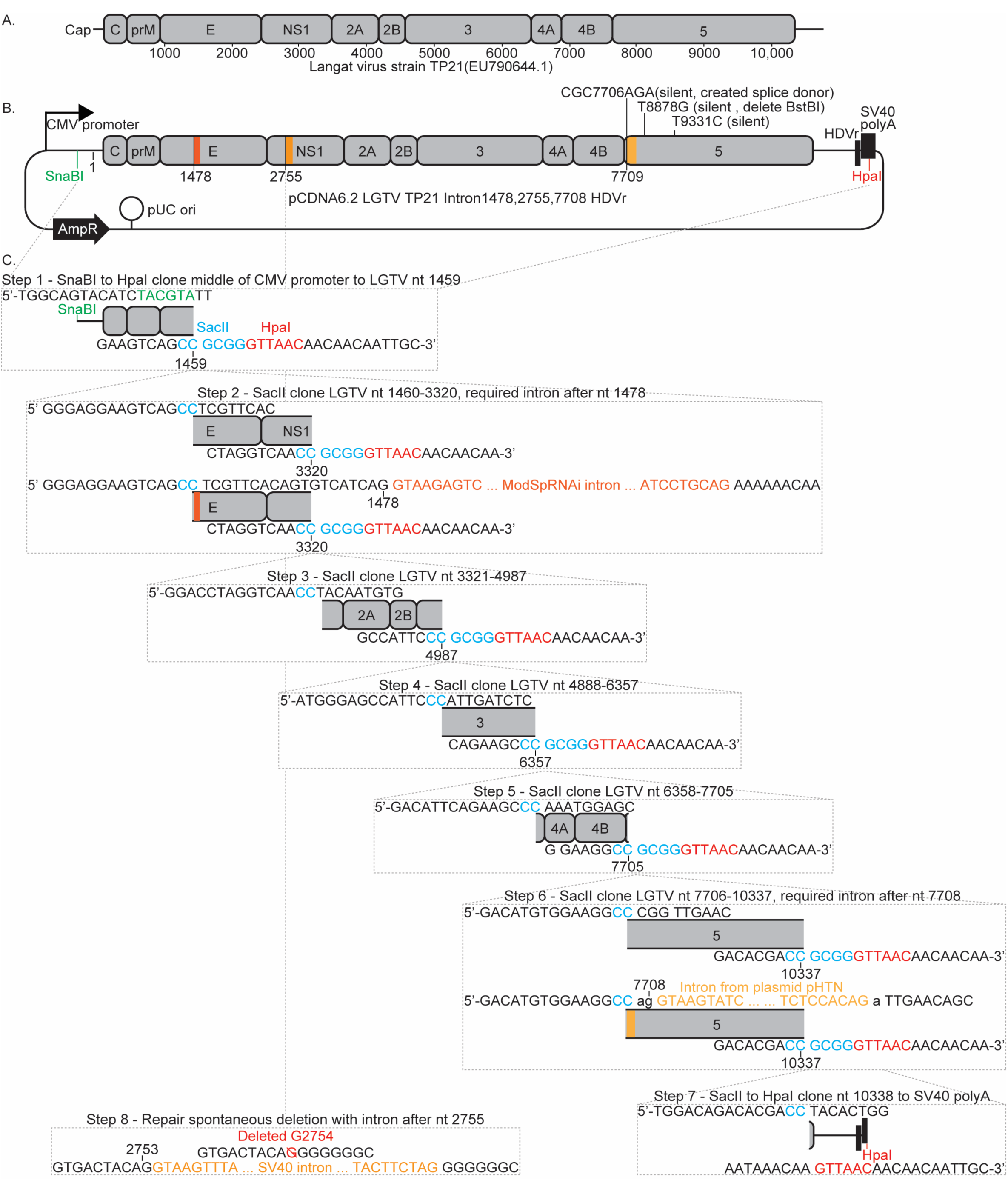
Illustrations of LGTV genome and cDNA plasmid. (A) Organization of the LGTV single stranded RNA genome showing the 5’ cap and positions of the mature viral proteins inside the single open reading frame. (B) Organization of the ‘pCDNA6.2 LGTV TP21 Intron1478,2755,7709 HDVr’ plasmid encoding the cDNA of the Langat virus strain TP21 (EU790644) genome under transcriptional control of the CMV promoter. A ribozyme (HDVr) is positioned to trim the RNA to have the LGTV 3’ end. Introns were inserted after nucleotides 1478, 2755, and 7709 of the viral sequence. The beta-lactamase resistance gene (AmpR), and origin of replication (pUC ori) are also marked. (C). Schematic representation of the steps used to create the above plasmid involved cloning synthetic DNA fragments to progressively assemble the entire genome. Each step graphically highlights the fragments added to the prior clone, and shows the sequence of the ends of each insert (see results and methods sections for details).

Both RNA- and DNA-launch systems are complicated by an instability of orthoflavivirus cDNA in bacteria, likely due to cryptic bacterial promoters that drive low-level expression of viral proteins and cause bacterial toxicity (reviewed in Aubry et al., 2015; Lindenbach, 2022). Several strategies have been developed to mitigate this problem, including the use of bacterial strains more tolerant of viral gene expression, propagation in very low copy-number plasmid backbones to minimize transcript levels, and the use of alternative hosts such as yeast. Another common approach is to divide the viral genome across multiple plasmids, each containing a portion of the cDNA that can be excised and ligated to reconstitute the full-length genome. The resulting complete cDNA can then serve as a template for *in vitro* transcription or, when placed downstream of a polymerase II promoter, for direct transfection into permissive cells.

An alternative strategy for stabilizing DNA-launch cDNA clones is to insert introns into regions that induce bacterial toxicity, thereby preventing translation in bacteria while allowing the authentic viral RNA sequence to be restored through splicing in mammalian cells. This approach has been successfully used to stabilize other positive-sense RNA viruses, including transmissible gastroenteritis coronavirus and Japanese encephalitis virus (González et al., 2002; Yamshchikov et al., 2001). We previously applied this strategy to generate a single-plasmid, CMV-driven infectious clone of the Zika virus MR766 isolate that could be stably maintained in *E. coli* and initiate infection directly in mammalian cells (Schwarz et al., 2016).

Here, we describe a high-copy, single-plasmid CMV/HDVr infectious cDNA clone of the LGTV TP21 strain that can be stably propagated in standard *E. coli* and produces high-titer infectious virus following direct DNA transfection of mammalian cells. A previous LGTV clone was stabilized by the addition of two introns, one in the NS1 region and another in NS5 (Tsetsarkin et al., 2016b). Building on this foundation, we employed an empirical strategy to identify and interrupt toxic regions, determining that three introns were required to fully stabilize the viral genome. Our study outlines this process in detail, providing a general roadmap for stabilizing flavivirus cDNA clones in bacteria. We further demonstrate the versatility of this system by generating fluorescent and luciferase protein–expressing reporter viruses, establishing practical tools for visualizing and quantifying LGTV infection. Together, these constructs provide a flexible, high-efficiency framework for engineering and studying tick-borne flaviviruses.

## Results

### Mapping and suppression of bacterial toxicity within the LGTV cDNA

We aimed to clone the full-length cDNA of the LGTV TP21 strain (Fig. 1A) into the high–copy-number plasmid pCDNA6.2 (Fig. 1B). In this construct, the authentic 5’ end of the viral sequence was positioned immediately downstream of the cytomegalovirus (CMV) promoter transcriptional start site, and the cDNA was followed by a hepatitis D virus ribozyme (HDVr) and the SV40 polyadenylation signal to produce transcripts with authentic 3’ termini of the viral genome. The resulting RNA is predicted to be identical to that of a genome delivered by an infectious virion and therefore capable of initiating infection following expression in host cells.

To empirically identify regions of the LGTV genome that are toxic in bacteria, we progressively cloned fragments of the viral cDNA starting from the 5’ end (scheme illustrated in Fig. 1C). We reasoned that fragments encompassing toxic regions would be difficult to clone, yielding few, slow-growing bacterial colonies. Furthermore, plasmids recovered from these colonies would likely contain nonsense or frameshift mutations that disrupt translation of downstream viral proteins responsible for the observed toxicity. The fragment spanning nucleotides 1460-3320 of the LGTV genome, corresponding to portions of the envelope (E) and nonstructural protein 1 (NS1) coding regions, was the first to display this phenotype (Fig. 1C, step 2). Only a single bacterial colony was obtained after transformation, and the plasmid recovered from this colony contained a two-nucleotide deletion at positions 1728-1729 that introduced a frameshift mutation. To alleviate this toxicity, we inserted an intron after nucleotide 1458, which allowed recovery of small but viable bacterial colonies that maintained the correct LGTV cDNA sequence without additional mutations.

The next three LGTV fragments (Fig. 1C, steps 3–5) were successfully cloned, although the resulting bacterial colonies continued to grow slowly. Attempts to clone a PCR fragment encoding the majority of the NS5 coding sequence, however, consistently failed to yield any colonies containing the insert (Fig. 1C, step 6). To overcome this, we introduced a second intron after nucleotide 7708 of the LGTV cDNA, modifying the adjacent CGG codon to a silent AGA mutation to generate consensus splice donor and acceptor sites. Whole-plasmid sequencing of the resulting full-length LGTV construct revealed a spontaneous single-nucleotide deletion at position 2754 (Fig. 1C, step 8), which likely arose during insertion of a downstream fragment and suggested residual toxicity associated with this region. To correct this, we added a third intron at the site of the deletion. The resulting three-intron plasmid could be stably propagated in *E. coli*, completing construction of a full-length LGTV cDNA clone under the transcriptional control of a CMV promoter. The final clone was identical to the reference LGTV TP21 sequence (GenBank accession NC_003690.1) except for the three introns, a silent mutation introduced to create splice donor and acceptor sites for the third intron (CGC7706AGA, numbering relative to the viral genome), two additional silent substitutions in NS5 (T8878G and T9331C), and one coding change in NS5, R542K (AGG9287AAG). The latter three mutations were carried over from the template plasmid used for NS5 amplification. The R542K substitution has been reported in multiple passaged LGTV isolates (Asghar et al., 2025; Campbell and Pletnev, 2000; Pletnev, 2001) and was therefore considered likely tolerated for viral replication.

### Cells transfected with the wild type LGTV plasmid produce infectious virus

To evaluate the production of infectious virus, we transfected 293T cells with the full-length, three-intron LGTV cDNA clone. Supernatants were collected daily from transfected cells and used to infect suitable target cells. Parallel cultures of Vero cells were infected with a patient-derived LGTV stock as a positive control, and mock-infected cells served as a negative control. Following infection, cultures were overlaid with a semisolid medium, and three days later, immunostaining with the 13A10 LGTV prM monoclonal antibody 4G2 revealed distinct foci of virus–positive cells in both the patient-derived virus and DNA-launched infections, but none in mock controls (Fig. 2A). The foci were larger in the patient-derived virus infection, suggesting that this virus spreads more efficiently in culture, which we quantified in subsequent growth-curve experiments.

**Figure 2.**
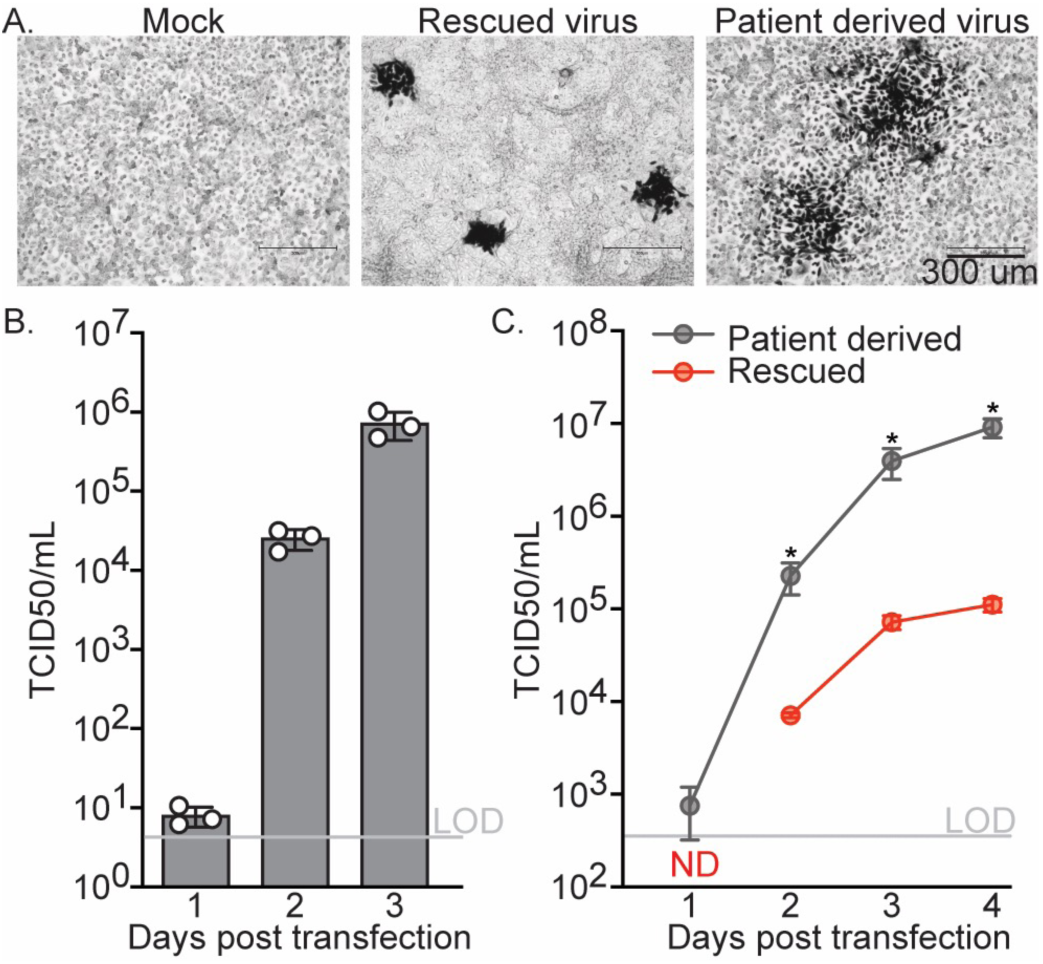
Characterization of plasmid rescued LGTV. (A) Representative 4G2 immunostaining of Vero cells infected with supernatants collected from 293T cells transfected with the wild-type LGTV plasmid, showing distinct foci of prM protein-positive cells. (B) Quantification of infectious virus released from transfected 293T cells over time. Supernatants collected at 1 to 3 days post-transfection were titered by TCID50 assay on Huh-7.5 cells, based on virus-induced cytopathic effect (CPE), and expressed as TCID50 per milliliter (TCID50/mL). Data represent the mean ± SEM of three independent transfections, each measured in triplicate. (C) Multicycle growth curves comparing the replication kinetics of the plasmid-rescued and patient-derived LGTV were performed by infecting Huh-7.5 cells at a multiplicity of infection (MOI) of 0.05 and measuring virus titers in culture supernatants collected daily by TCID50 assay. Titers are expressed as TCID50/mL. Data represent means ± SEM of three independent experiments, each performed in triplicate. *P < 0.05 (unpaired t-test with Welch’s correction).

Infectious virus produced from transfected 293T cells was quantified by endpoint dilution assay on Huh-7.5 cells (TCID50), which are highly permissive to flavivirus infection due to a dysfunctional RIG-I gene (Blight et al., 2002), based on LGTV-associated cytopathic effect (CPE). At 24 hours post-transfection, supernatants from 293T cells transfected with the LGTV plasmid contained 5.6x10^0^ TCID50/mL of infectious virus, which increased to 1.63x10^4^ and 5.0x10^5^ TCID50/mL at two and three days post-transfection, respectively (Fig. 4B). These titers were somewhat lower than the 10^7^ to 10^8^ TCID50/mL peak titers typically observed for patient-derived or laboratory-passaged LGTV stocks (Tsetsarkin et al., 2016b), likely reflecting the limited permissiveness of 293T cells to flavivirus infection. These cells are primarily used for their high transfection efficiency rather than for productive viral replication.

### Comparison of the growth properties of rescued and patient-derived LGTV

To directly compare the growth properties of plasmid-rescued LGTV with the patient-derived inoculum, multicycle growth curve experiments were performed in which Huh-7.5 cells were infected at a multiplicity of infection (MOI) of 0.05 with either the rescued wild-type or patient-derived virus. Supernatants were collected daily for four days post-infection and titrated by TCID50 assay on Huh-7.5 cells. The resulting growth curves revealed that the rescued virus replicated more slowly than the patient-derived virus (Fig. 2C). At one day post-infection, virus released from patient-derived infections was just detectable, whereas virus from rescued infections remained below the limit of detection. At all subsequent time points, the patient-derived virus produced 32- to 82-fold higher titers than the rescued virus. These results parallel previously reported multicycle growth curves for rescued 2010 Cambodia and 2015 Brazil Zika virus strains, which displayed approximately 10- to 50-fold slower spread compared to their respective patient-derived isolates (16, 19). Also, as described above, our rescued LGTV clone differs from the published TP21 sequence by several noncoding and coding changes. Although the exact sequence of our patient-derived isolate is unknown, such sequence differences may contribute to the observed attenuation.

### Construction of an LGTV fluorescent protein reporter virus

To enable visualization of LGTV infection, we engineered a reporter virus expressing the red fluorescent protein mCherry. The design followed a widely used orthoflavivirus reporter format in which the reporter gene is inserted as a translational fusion at the amino terminus of the viral polyprotein. To preserve essential RNA elements, we retained the first 28 codons of the capsid (C) protein upstream of the reporter gene, maintaining sequences that form secondary structures required for viral RNA replication through base pairing with the 3’ untranslated region. Downstream of the reporter, we introduced the foot-and-mouth disease virus (FMDV) 2A peptide to mediate co-translational cleavage and reinitiation of translation at the authentic start codon of the full-length capsid protein. The first 28 codons of the full-length C gene were recoded with silent mutations to prevent complementarity with the upstream C28 sequence, thereby preserving the essential RNA secondary structures. We first constructed an “acceptor” plasmid containing the duplicated C28 sequence (with silent mutations) followed by the FMDV 2A peptide (Fig. 3A). A unique SacII restriction site was introduced between the wild-type C28 sequence and the FMDV 2A peptide to permit insertion of reporter genes. The mCherry coding sequence was then cloned into this site to generate the mCherry reporter plasmid (Fig. 3B).

**Figure 3.**
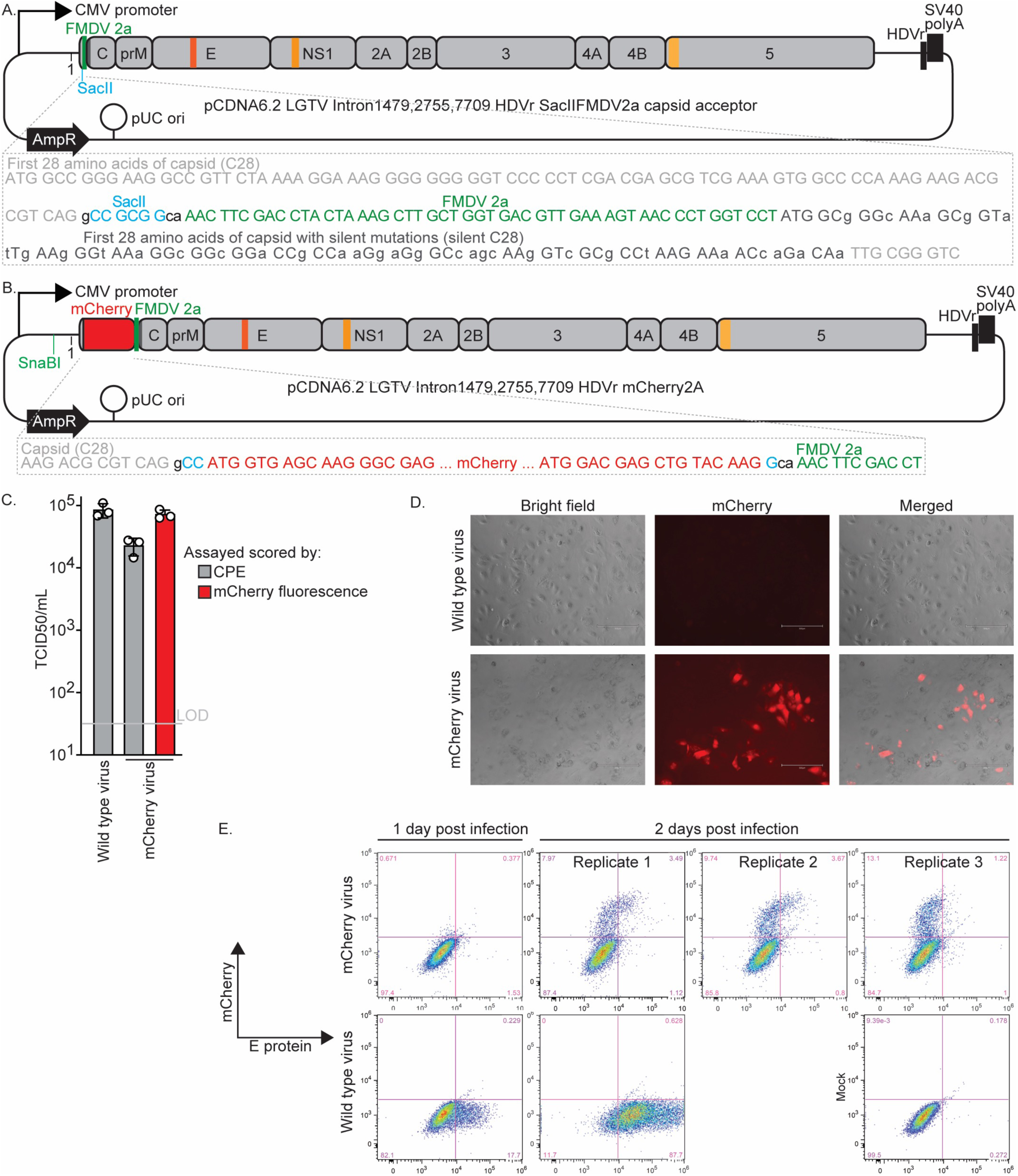
Characterization of fluorescent reporter protein-expressing LGTV. (A) Schematic of the ‘acceptor plasmid’ subsequently used for insertion of reporter genes at a unique SacII site as translational fusions to the first 28 amino acids of the capsid (C) protein and followed by the FMDV 2A peptide and the full-length polyprotein bearing silent mutations in the duplicated C28 region. The nucleotide sequence of this region are shown below for clarity. (B) Schematic of the mCherry fluorescent protein–expressing LGTV plasmid, with the sequence of the insertion shown for clarity. (C) Rescue titers at 3 day post transfection of wild-type (non-reporter) and mCherry-expressing LGTV. Titers were determined by TCID50 assay on Huh-7.5 cells based on cytopathic effect (CPE); the mCherry virus was additionally scored by fluorescence. Data represent means ± SEM of three independent experiments, each performed in triplicate. (D) Representative fluorescence and phase-contrast images of Huh-7.5 cells infected with the indicated viruses, 3 days post-infection. (E) Representative flow cytometry plots of Huh-7.5 cells infected with the indicated viruses at 1 or 2 days post-infection. The x-axis shows E protein staining detected with the 4G2 antibody, and the y-axis indicates mCherry fluorescence. The right panels show three independent replicate infections with independently produced mCherry virus stocks at 3 days post-infection. Mock-infected cells are included as controls.

Transfection of 293T cells with this plasmid resulted in the production of infectious virus that induced both cytopathic effect (CPE) and mCherry fluorescence in Huh-7.5 cells (Fig. 3C). The rescue titer of this virus was 2.28x10⁴ TCID50/mL at three days post-transfection, approximately 29-fold lower than that of the nonreporter wild-type virus (Fig. 3D). Such attenuation is typical for reporter orthoflaviviruses (Lindenbach, 2022). When TCID50 assay wells were scored for mCherry fluorescence by microscopy, the calculated titers were 3.7-fold higher than those determined by CPE, suggesting that the fluorescent reporter provides a more sensitive measure of infection.

To further assess the kinetics and stability of mCherry expression, Huh-7.5 cells were infected with the reporter virus and analyzed by flow cytometry at multiple time points. Fluorescence was barely detectable at one day post-infection but became robust by two days post-infection (Fig. 3E). Quantitatively, mCherry fluorescence provided a 3.5-fold more sensitive readout of infection than E-protein immunostaining with the 4G2 monoclonal antibody, as determined by dividing the number of all mCherry-positive events by the number of all 4G2-positive events across the three replicates shown in Fig. 3E. The FACS analysis also confirmed that the virus stock produced by plasmid transfection exhibited no detectable loss of the reporter gene, a common occurrence for orthoflavivirus insertions that reduce fitness. Such loss would manifest as cells strongly positive for E protein but negative for mCherry fluorescence, which would appear in the lower-right quadrant of the mCherry virus replicates in Fig. 3E, but was not observed.

### Construction of an LGTV luciferase protein reporter virus

To develop a quantitative and sensitive assay for LGTV infection, we used the SacII-FMDV 2A-capsid acceptor plasmid to generate a virus encoding the Gaussia luciferase (GLuc) gene (Fig. 4A). Transfection of this plasmid into 293T cells yielded infectious virus titers approximately 89-fold lower than those obtained with the wild-type, nonreporter construct, as determined by CPE-based TCID50 assay on Huh-7.5 cells (Fig. 4B). Such attenuation is typical of reporter orthoflaviviruses and can occasionally promote loss of the reporter through recombination. Infection of Huh-7.5 cells with the rescued GLuc virus produced a dose- and time-dependent increase in luciferase activity (Fig. 4C). Strong luminescence was detectable as early as one day post-infection, and the magnitude of this signal correlated with the inoculum dose. Continued incubation resulted in further increases in GLuc activity, consistent with either progressive viral replication in initially infected cells or subsequent spread to naïve cells within the culture.

**Figure 4.**
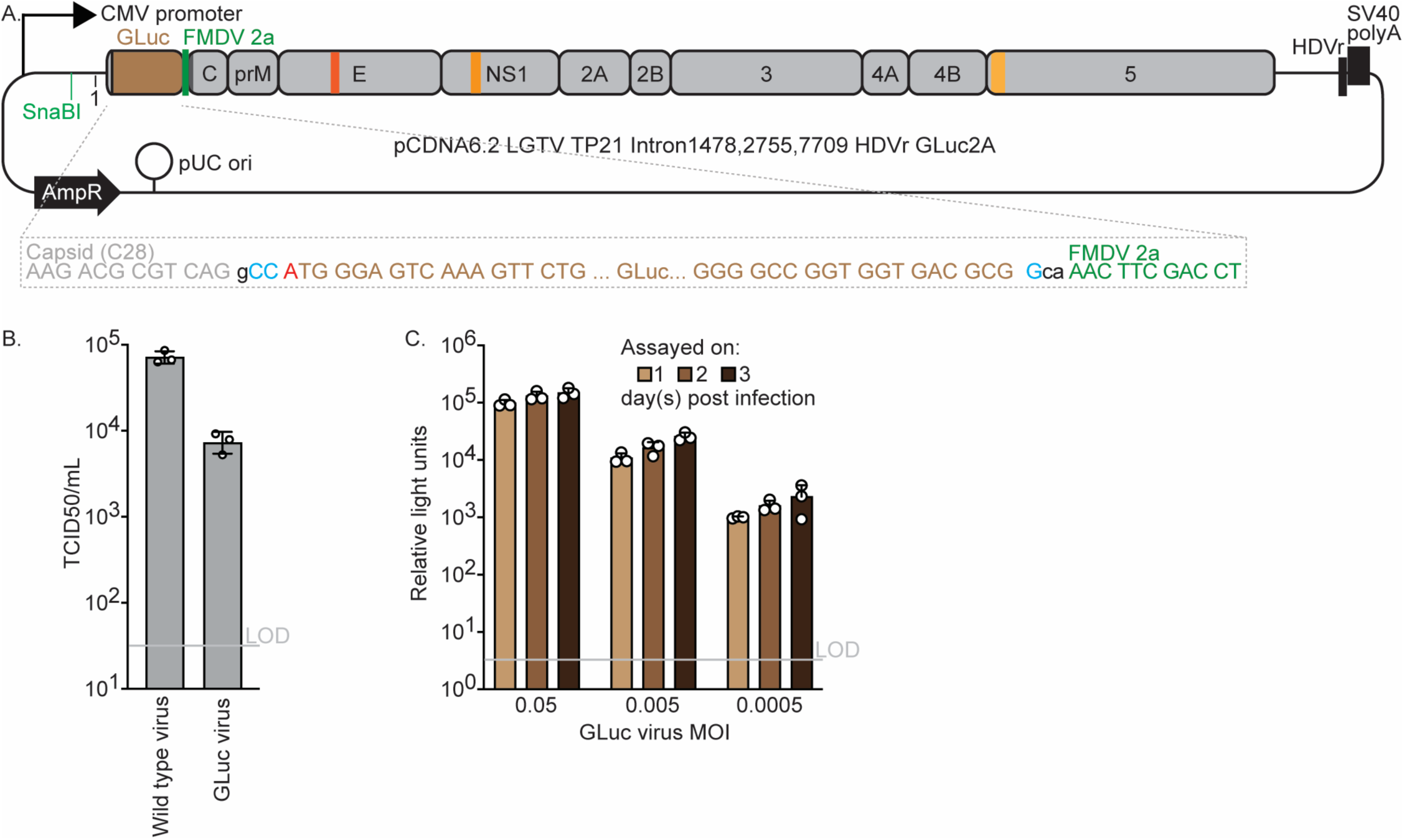
Characterization of luciferase reporter protein-expressing LGTV. (A) Schematic of the Gaussia luciferase (GLuc) reporter LGTV plasmid, showing the GLuc insertion site and sequence context. (B) Rescue titers at 3 day post transfection of wild-type (non-reporter) and GLuc-expressing LGTV. Titers were determined by TCID50 assay on Huh-7.5 cells based on cytopathic effect (CPE). Data represent means ± SEM of three independent experiments, each performed in triplicate. (C) Luciferase activity in Huh-7.5 cells infected at the indicated multiplicities of infection (MOI = infectious units per cell) and assayed at 1, 2, and 3 days post-infection. Data represent means ± SEM of three independent experiments, each performed in triplicate.

## Discussion

The goal of this study was to develop a simple and robust reverse-genetics system for LGTV, a tick-borne flavivirus that serves as a biosafety level 2 model for TBEV. We constructed a single-plasmid, CMV promoter-driven infectious cDNA clone based on the LGTV TP21 strain that can be stably propagated in standard *E. coli* and efficiently launches infectious virus upon transfection of mammalian cells. The resulting plasmid incorporates three introns that interrupt regions of the viral genome that induce toxicity in bacteria. Using this clone, we generated both fluorescent and luciferase reporter derivatives that facilitate quantitative and high-throughput analyses of infection.

Our findings build upon the work of Tsetsarkin and colleagues (2016b), who first generated a CMV-driven, intron-stabilized LGTV clone by introducing two introns, with one in NS1 and another in NS5, to stabilize the viral cDNA in bacteria and enable recovery of infectious virus (Tsetsarkin et al., 2016b). Similarly, our construct required introns in NS1 and NS5, and inclusion of a third intron within the E coding region further improved plasmid stability by reducing residual toxicity. Our present work complements that of Tsetsarkin and colleagues by detailing an empirical approach to identify and interrupt bacterial toxicity across the LGTV genome. This stepwise mapping of toxic regions provides a generalizable framework for stabilizing flavivirus cDNA clones and highlights that the number and location of required introns can vary depending on plasmid context and promoter configuration. Two design features were particularly important to our strategy. First, although large DNA fragments (>5 kb) can now be readily synthesized, we used smaller fragments (<2 kb) to provide finer resolution of toxic regions. Second, we cloned sequentially from the 5’ to the 3’ end of the genome, reasoning that cryptic bacterial promoters upstream of the viral sequence could drive expression of downstream viral proteins and thereby cause toxicity. Adding introns at the 5’ boundary of fragments that exhibited toxicity proved an efficient and effective means to disrupt these cryptic transcripts and stabilize the full-length viral cDNA.

Our DNA-launched LGTV clone produced high levels of infectious virus in mammalian cells, confirming that transcription from the CMV promoter yields replication-competent RNA capable of initiating the full viral life cycle. Following transfection of 293T cells, virus production increased steadily over three days, reaching titers of approximately 5x10⁵ TCID50/mL. Infection of highly permissive Vero and Huh-7.5 cells demonstrated that the DNA-launched virus produced characteristic cytopathic effect and spread efficiently in culture, validating the functionality of the cloned genome.

In multicycle growth-curve experiments, however, the rescued LGTV replicated more slowly than the patient-derived isolate, producing 32- to 82-fold lower titers at later time points. Similar patterns of modest attenuation have been reported for DNA-launched Zika viruses, suggesting that this is a common phenotype of plasmid rescued flaviviruses (Shan et al., 2016; Tsetsarkin et al., 2016a). The reduced replication kinetics may reflect the highly genetically homogenous virus population, while the passaged patient-derived virus has built-in sequence diversity which may be beneficial to the capacity to replicate in diverse conditions (Asghar et al., 2016; Helmová et al., 2020; Litov et al., 2018). Alternatively, sequence differences between our cloned TP21 genome and the patient-derived LGTV stock could contribute to the observed attenuation. The rescued clone contains several synonymous substitutions and one amino-acid change (R542K) in NS5. Notably, this R542K substitution has been reported in multiple LGTV isolates (Asghar et al., 2025; Campbell and Pletnev, 2000; Pletnev, 2001), indicating that it is tolerated, but effects on replication cannot be excluded. Despite this attenuation, the rescued LGTV retained robust infectivity and spread in cell culture, supporting its use for reverse-genetics applications. Indeed, this system provides an easy launching point for further investigations into LGTV replication and attenuation phenotypes.

The reporter derivatives expand the versatility of the LGTV infectious clone. The mCherry virus enables direct visualization of infection and spread, whereas the secreted Gaussia luciferase (GLuc) reporter provides a sensitive, quantitative, and non-destructive measure of replication. Both reporters were introduced using a modular acceptor plasmid design that allows straightforward exchange of fluorescent or luminescent genes for alternative applications. To our knowledge, no fluorescent protein–expressing LGTV has previously been reported. However, a related luciferase-expressing LGTV clone encoding NanoLuc in a low–copy-number bacterial plasmid was recently described (Miao et al., 2024). Although both of the reporter viruses generated in this study exhibited lower titers than the nonreporter clone, such attenuation is typical of orthoflavivirus reporters and likely reflects the metabolic cost of reporter expression and the increased genome length. Importantly, both of our reporter viruses retained stability following rescue, with no evidence of recombination-mediated loss during early passages.

In summary, we have developed a stable, high-copy, CMV-driven LGTV infectious clone and accompanying reporter viruses that enable direct DNA launch of infection in mammalian cells. This system overcomes obstacles to flavivirus genome stability in bacteria and provides a versatile toolkit for dissecting viral replication, immune evasion, and pathogenesis. By combining genetic precision with biosafety-level-2 accessibility, these constructs establish LGTV as a practical model for tick-borne flavivirus biology and a foundation for future vaccine and antiviral development.

## Materials and Methods

### Cell lines and culture

293T and Huh-7.5 cells (provided by Charles M. Rice, Rockefeller University, New York, NY) were grown as previously described (Hopcraft et al., 2016) in Dulbecco’s Modified Eagle’s Medium (DMEM; Gibco BRL Life Technologies, Gaithersburg, MD) with 10% fetal bovine serum (FBS; Gibco BRL Life Technologies, Gaithersburg, MD), and 108.8 units/mL penicillin and 108.8 µg/mL streptomycin (Gibco BRL Life Technologies, Gaithersburg, MD).

### LGTV stock propagation and infectious virus quantification

The LGTV TP21 patient derived virus stock was obtained from Alexander Pletnev (NIH, NIAID) and was propagated in Vero and Huh-7.5 cells (Campbell and Pletnev, 2000). To amplify the patient derived virus stock, 1x10^4^ Huh-7.5 cells were seeded in a 10cm petri dish one day prior to infection. The dish was infected with a MOI of 0.5 Huh-7.5 cell TCID50 infectious units per cell in DMEM with 3% FBS, 108.8 units/mL penicillin, and 108.8 µg/mL streptomycin. One day post infection, the supernatant was aspirated and replaced with 8 mL DMEM with 3% FBS, 108.8 units/mL penicillin, and 108.8 µg/mL streptomycin. Two and three days post infection, the supernatant was collected and clarified by passage through 0.45-μm filters. The three day post infection supernatant was used in all experiments that tested the patient derived virus.

Virus titers were determined by CPE-based limited dilution assay on Huh-7.5 cells, following established protocols (Fulton et al., 2017). Here, 1x10^4^ cells were seeded in each well of a 96-well plate the day before being infected. For most infections, we used 50 µLof virus serially diluted in DMEM with 10% FBS, 108.8 units/mL penicillin, and 108.8 µg/mL streptomycin (8 wells per dilution). For the one day post transfection supernatants, we used 87 µL per well to maximize sensitivity. Wells were scored for LGTV-induced CPE, which was observed as early as 5 days post transfection, but final scoring was conducted on day 7, and infectious titers were calculated as TCID50/mL according to the method of Reed and Muench (Reed and Muench, 1938).

### Plasmid construction

Our prior ‘pCDNA6.2 Zika MR766 Intron3115 HDVr’ plasmid encoding the ZIKV MR766 isolate cDNA under the control of the CMV promoter (Schwarz et al., 2016) served as the backbone for construction of the LGTV cDNA clone. The ZIKV cDNA sequence was replaced with the TP21 strain LGTV cDNA (GenBank accession no. NC_003690.1), with the 5’ and 3’ termini precisely exchanged to maintain authentic viral boundaries. Our final plasmid encoded four sequence changes in the LGTV TP21 cDNA from the reference sequence (GenBank accession no. NC_003690.1): three silent mutations, CGC7706AGA (introduced to create splice donor and acceptor sites for the third intron), T8878G, and T9331C—and one coding change, AGG9287AAG, resulting in an R542K amino acid substitution in NS5.

We used a multistage cloning strategy to assemble the full length LGTV cDNA clone (illustrated in Fig. 1C). The first fragment, a 1,739-nt synthetic DNA (Twist Bioscience, San Francisco, CA), contained sufficient overlap with the SnaBI restriction site in the CMV promoter of pCDNA6.2 to permit In-Fusion cloning (Takara Bio USA, Inc., San Jose, CA). This fragment extended through nucleotide 1459 of the LGTV genome, ending in two consecutive cytosines (C1458-1459), which were modified to create a unique SacII restriction site. The downstream end of the fragment included overlap with the vector’s HpaI site located adjacent to the SV40 polyadenylation signal. Using the In-Fusion HD Cloning Kit, this fragment was inserted between the SnaBI and HpaI sites of our previously described pCDNA6.2-ZIKV-MR766-Intron3115-HDVr plasmid (Fig. 1C, step 1) (Schwarz et al., 2016).

Subsequent synthesized LGTV cDNA fragments were sequentially introduced by In-Fusion cloning into the SacII site at the 3’ end of each preceding insert. This approach preserved only the two terminal cytosines of the prior fragment while replacing the remainder of the SacII site with authentic LGTV sequence. In-Fusion cloning thereby enabled us to use transient, uniquely placed SacII sites as flexible junctions between fragments, each created at any desired location containing two adjacent cytosines, allowing precise, seamless assembly of the complete LGTV cDNA. The NS5 sequence cloned in step 6 of Figure 1, was PCR amplified from a pCAGG-LGTV plasmid obtained for the laboratory of Adolfo Garcia-Sastre (Icahn School of Medicine at Mount Sinai, New York, NY).

Plasmids were propagated in either NEB Turbo Competent *E. coli* (New England Biolabs, Ipswich, MA) that grow rapidly and thus compensate for flavivirus cDNA induced toxicity, or Stellar cells (Takara Bio USA, Inc., San Jose, CA). These bacteria were grown at 37°C for all cloning and amplification steps, including recovery following heat shock transformation.

To alleviate bacterial toxicity associated with the LGTV genomic region spanning nucleotides 1459-3320, the ModSpRNAi intron (Ying et al., 2010) was added after LGTV nucleotide 1478. This site which bears the CAG and A flanking sequences contains the canonical exon sides of splice donor (C or A, A, and G) and acceptor motifs (G or A) required for efficient splicing in mammalian cells. Toxicity within the NS5 coding region was mitigated by introducing the synthetic intron from plasmid pHTN (Promega, Madison, WI; GenBank accession no. JF920304) after nucleotide 7709. This insertion required the inclusion of silent mutations (CGC7706AGA) to generate functional splice donor and acceptor sequences. Finally, the toxicity indicated by a spontaneous deletion of nucleotide G2754 was alleviated by inserting the SV40 intron from pCMV4 (GenBank accession no. AF239248)(Andersson et al., 1989) after LGTV nucleotide 2755. We used three different introns to avoid incorporating repeated sequences throughout the plasmid that can lead to instability.

### 293T cell transfection and supernatant collection

Plasmid-based LGTV was produced in 293T cells by transfections performed using techniques described previously (Fulton et al., 2017). One day prior to transfection, 5x10^5^ cells/well were seeded in 6-well poly-lysine coated plates. Cells were transfected with 2 μg of DNA per well, with 100 μL of Opti-MEM (Gibco BRL Life Technologies, Gaithersburg, MD) and 6 μL of TransILT (Mirus Bio, Madison, WI) that had been incubated with the DNA for 20 to 60 minutes prior to transfection. Supernatants were collected daily from one to three days post-transfection and clarified by passage through 0.45 μm filters.

### Focus forming assay

To visualize infected cells by focus forming assay, 150 µL of virus-containing supernatant was adsorbed on Vero cell monolayers for one hour at 37 °C in 24-well plates. Cells were then overlaid with 1 mL of DMEM (Gibco BRL Life Technologies, Gaithersburg, MD) supplemented with 0.8% methyl cellulose, 2% FBS, 100 units/mL penicillin, and 100 µg/mL streptomycin. Cells were incubated for two days at 37°C and then washed with PBS (Gibco BRL Life Technologies, Gaithersburg, MD), followed by fixation with 100% methanol overnight.

After washing with PBS, plates were incubated for 1 h at 37°C with 500 µL of a 1:1000 dilution of LGTV-13A10 antibody (BEI Resources, Manassas, VA). After three washes, 250 µL per well of goat anti-mouse horseradish peroxidase labelled antibody (4 µg/mL; ThermoFisher Scientific, Waltham, MA) was added and incubated for one hour at 37 °C, followed by three additional PBS washes. LGTV foci were visualized by the addition of 500 µL of DAB mix, prepared by dissolving 450 µg/mL DAB (ThermoFisher Scientific, Waltham, MA) and adding 0.45 µL/mL of 30% H₂O₂ (ThermoFisher Scientific, Waltham, MA) in PBS. The reaction was carried out at room temperature for 10 min and stopped by washing with water.

### Growth curves

One day prior to infection, 1x10^5^ Huh-7.5 cells per well were seeded in a 24-well poly-lysine coated plate in triplicate. Then, the cells were infected at a MOI of 0.05 Huh-7.5 cell TCID50 infectious units per cell in DMEM with 3% FBS, 108.8 units/mL penicillin, and 108.8 µg/mL streptomycin. Supernatants were collected daily for the next four days. Supernatant infectivity was assayed as previously described.

### Fluorescence microscopy

Bright field and fluorescence images were acquired with the EVOS M5000 microscope (ThermoFisher Scientific, Waltham, MA).

### Flow cytometry

Cells were fixed with 1% paraformaldehyde and stained with the primary 4G2 flavivirus E antibody (kind gift from Florian Krammer, Icahn School of Medicine at Mount Sinai, New York, NY) and a goat anti-mouse secondary antibody conjugated to Alexa fluorophore 647 (ThermoFisher Scientific, Waltham, MA), using methods previously described (Veit et al., 2024). Cells were analyzed using an Attune NxY flow cytometer (ThermoFisher Scientific, Waltham, MA).

### Luciferase assay

One day prior to infection, 5x10^4^ Huh-7.5 cells were seeded in a 24 well poly-lysine coated plate. Cells were infected with 0.05, 0.005, 0.0005 infectious units per cell, each in triplicate. Cells were lysed one, two, and three days post infection with Renilla Luciferase Assay Lysis Buffer (Promega, Madison, WI) and stored at -80 °C. For determination of luciferase activity, lysates were thawed and 10 µL of each were assayed with 40 µL of substrate from the Renilla Luciferase Assay System (Promega, Madison, WI) and quantified with a BioTex Synergy 4 multidetection microplate reader (BioTek, Winooski, VT).

### Statistical analysis

Data were analyzed using an unpaired t-test with Welch’s correction, \**P*<0.05, \*\**P*<0.005, \*\*\**P*<0.0005 on Prism software (GraphPad Software). Values in graphs represent the mean and standard error of experiments performed in triplicate or quadruplicate, with between two and five independent experiments.

## Acknowledgements

The authors thank Charles Rice (Rockefeller University, New York, NY) for 293T and Huh-7.5 cells, Adolfo Garci-Sastre for the pCAGGS-LGTV NS5 plasmid (Icahn School of Medicine at Mount Sinai, New York, NY), Alexander Pletnev for the LGTV TP21 clinical isolate (NIAID/National Institutes of Health), and Florian Krammer for 4G2 monoclonal antibody (Icahn School of Medicine at Mount Sinai, New York, NY). We would like to thank the expertise and assistance of the Dean’s Flow Cytometry CORE at Mount Sinai.

